# Current strategic limitations of phylogenetic tools badly impact the inference of an evolutionary tree

**DOI:** 10.1101/2021.01.21.427545

**Authors:** Shamantha Nasika, Ashish Runthala

## Abstract

For drawing an evolutionary relationship among several protein sequences, the phylogenetic tree is usually constructed through maximum likelihood-based algorithms. To improve the accuracy of these methodologies, many parameters like bootstrap methods, correlation coefficient and residue-substitution models are presumably over-ranked to derive biologically credible relationships. Although the accuracy of protein sequence alignment and the substitution matrix are preliminary constraints to define the biological accuracy of the overlapped sequences/residues, the alignment is not iteratively optimized through the statistical testing of residue-substitution models. The study majorly highlights the potential pitfalls that significantly affect the accuracy of an evolutionary protocol. It emphasizes the need for a more accurate scrutiny of the entire phylogenetic methodology. The need of iterative optimizations is illustrated to construct a biologically credible and not mathematically optimal tree for a sequence dataset.

## Introduction

An evolutionary relationship is usually drawn to screen the sequence/functional similarity among the protein sequences (Chan 1989; Kamjula et al. 2020; Keeling et al. 1998; Sharma et al. 2018; T. Lanisňik Rizňer 2001). On the basis of mutual sequence similarity, the phylogenetic tree is constructed as the schematic depiction of the evolutionary connection between the species of different origins, and evolutionary distance/relatedness is assessed through several non-absolute scores including branch length, maximum likelihood score, and topology.

To draw the evolutionary relationships, a sequence profile or multiple sequence alignment (*MSA*) is usually constructed through several protocols, viz. MAFFT, MUSCLE, Kalign, TCoffee, Clustal-Omega and PROMALS3D(Di Tommaso et al. 2011; Edgar 2004; Kazutaka Katoh 2002; Lassmann and Sonnhammer 2005; Pei and Grishin 2014; Sievers et al. 2011). Multiple sequence comparison by log-expectation (MUSCLE; https://www.ebi.ac.uk/Tools/msa/muscle/) and multiple alignments using fast fourier transform (MAFFT; https://www.ebi.ac.uk/Tools/msa/mafft/) protocols are better than ClustalW and quickly align hundreds of sequences within seconds (Edgar 2004; Kazutaka Katoh 2002). Kalign alignment is also a very fast MSA tool which is applicable for large alignments. It uses WU-Manber string-matching algorithm to quickly align the sequences (Kazutaka Katoh 2002; Lassmann and Sonnhammer 2005). However, the profile multiple alignment with predicted local structures and topological constraints (PROMALS3D) proficiently aligns sequences/structures on the basis of sequence profiles, predicted secondary structures and structural constraints (http://prodata.swmed.edu/promals3d/promals3d.php). In contrast to these servers, Clustal-Omega is capable of aligning numerous sequences on the basis of a guide tree (Sievers et al. 2011) and hidden markov model (*HMM*) profiles (Sievers and Higgins 2018). Residue substitution matrix is used to align the sequences, and is deployed by the phylogeny protocol for deriving the evolutionary relationships. Phylogeny is routinely derived through nine different substitution matrices, viz. Dayhoff (Winona C. Barker 1978), JJT, BLOSUM62, WAG, PMB, DCMut (Kosiol and Goldman 2005), JTTDCmut (Kosiol and Goldman 2005), LG (Le and Gascuel 2008), and VT (Alva et al. 2016).

On the basis of sequence alignment, evolutionary relationship is usually drawn through several tools including a tree and reticulogram reconstruction (*T-REX),* MEGA5, PhyML, randomized axelerated maximum likelihood (RAxML), NGPhylogeny.fr and IQTree. PhyML uses the maximum likelihood method along with bootstrap and other parameters (Guindon et al. 2005). While T-REX deploys maximum likelihood/parsimony methods (Boc et al. 2012), MEGA5 and NGPhylogeny.fr offer a customizable platform (Hall 2013; Lemoine et al. 2019). However, the RAxML protocol is routinely used for large sequence datasets (Stamatakis 2014). In comparison to these strategies, IQTree (Nguyen et al. 2015) offers a highly customizable protocol to build even the complex trees more accurately. The method uses the ultrafast bootstrap algorithm and saves the runtime through statistical scoring, and is significantly faster than RAxML and PhyML (Hoang et al. 2018; Minh et al. 2013).

Evolutionary analysis has been extensively used for various research projects since 1883, and until December 18, 2019, 207863 articles are published, ~1540 articles every year (BROOKS 1883; Xuan et al. 2019). Almost all phylogenetic algorithms are maximally dependent on the aforementioned parameters and the impact of a biologically inaccurate sequence alignment and an incompatible residue substitution matrix is greatly disregarded. Most of these methods deploy a default substitution/alignment protocol for every sequence dataset (Boc et al. 2012; Guindon et al. 2005; Hall 2013; Lemoine et al. 2019; Nguyen et al. 2015; Stamatakis 2014), and do not train the dataset over available options. Moreover, no previous study has simultaneous assessed the quantified the negative impact of an incorrect substitution matrix, alignment protocol and the depth of a sequence dataset over the accuracy of a phylogenetic tree. By providing a comprehensive assessment of all the alignments and substitution models over an evolutionary tree, the study proves the importance of well-trained substitution model and biologically correct alignment. To aid the construction of a more accurate evolutionary server, the study defines the vital constraints for a heuristic and iterative tailoring of input data. It will aid us extract the closest template structures for consistently predicting the near-native protein models over the conventional protocols (Ashish and Shibasish 2013; Runthala 2012; Runthala 2015; Runthala 2020; Runthala and Chowdhury 2014; Runthala and Chowdhury 2016; Runthala and Chowdhury 2019), and will allow us extract a more-accurate functional relationship among the candidate sequences for various research domains (Amrein et al. 2019; Anil Kumar Jamithireddy 2020; Ashish Runthala 2011; Phulara et al. 2020).

## Materials and methods

### Building the sequence dataset and evaluating the sequence identity

For rigorously doing the evolutionary analysis though several parameters and to save the computational time, the families with less than 1000 sequences are prioritized. As the accuracy of a phylogeny protocol is strongly dependent on the number of sequences and the sequence length, three non-redundant datasets 1-3, containing 36, 109 and 712 sequences, are orderly derived from three randomly-selected families PF18258, PF18760 and PF00074 (Figure 1). As simulating the sequence datasets on basis of various constraints might lead to a biased perturbation of their individual and mutual diversity, the three arbitrarily defined datasets should be robustly accurate and more fruitful for the study. As sequence identity is an important basis of evolutionarily clustering the sequences for estimating their sequence divergence, the mutual identity matrix is computed for each of these datasets through the Clustal-omega protocol (Sievers et al. 2011).

**Figure 1:**
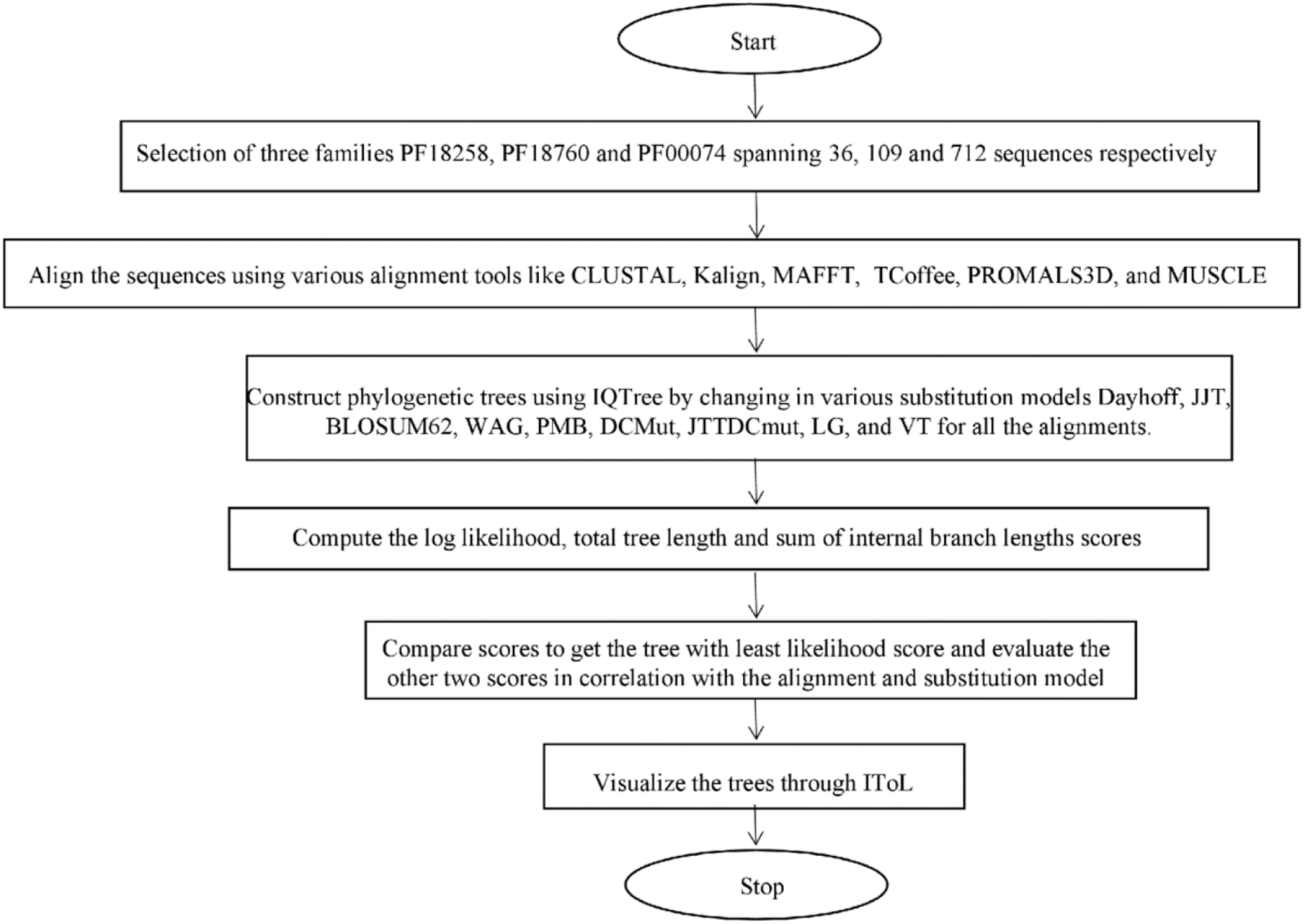
Flowchart representation of the methodology deploying the construction of three varied sequence datasets and the evolutionary analysis of their several alignments through diverse substitution models.

### Construction of phylogenetic trees

The constructed sequence datasets are aligned through Clustal-Omega, Kalign, MAFFT, MUSCLE, TCoffee and Promals3D, and the HMM-based Clustal-Omega (*Clustal1*), Kalign (*Kalign1*), MAFFT (*MAFFT1*), MUSCLE (*MUSCLE1*) and TCoffee (*TCoffee1*) protocols (Alva et al. 2016) to screen the more accurate alignment on the basis of phylogenetic tree. For the computational limit of 2500-character length of an input alignment, TCoffee protocol for both EBI and HHPred servers could not be deployed to align the set2 and set3 datasets. Likewise, for the sequence-strength limit of 500 entries, the MUSCLE alignment of EBI server is excluded for set3. A set of nine substitution models viz. Dayhoff (Winona C. Barker 1978), JJT, BLOSUM62, WAG, PMB, DCMut (Kosiol et al. 2006; Kosiol and Goldman 2005), JTTDCmut (Kosiol et al. 2006; Kosiol and Goldman 2005), LG(Le and Gascuel 2008), and VT (Alva et al. 2016) are likewise screened to decipher the more compatible matrix for a sequence dataset.

To quickly construct the evolutionary tree through an ultrafast bootstrap methodology on the basis of a customized protocol, the IQ-Tree server is used (Nguyen et al. 2015). The maximum likelihood based phenetic analysis parametrically estimates the evolutionary tree through several features including branch length, nucleotide composition bias and corrected distances between taxa (Nabhan and Sarkar 2012). As the theoretical probability is dependent on the observed data and not the experimental results, the correlation coefficient is further evaluated to find more accurate phylogenetic solutions (Bruce Rannala 1996; Kensche et al. 2008; Nunes et al. 2015). For each of the constructed sequence alignments, the evolutionary trees are derived through IQTree at the correlation coefficient of 0.9. As bootstrapping assesses the robustness of a phylogenetic solution (Felsenstein 1985), a quick 1000-bootstrap run is deployed to decipher the biologically-feasible substitution matrix and alignment protocol for each of the sequence datasets.

### Evolutionary analysis

The maximum-likelihood based tree is assessed through log-likelihood, total tree length and the total internal branch length score. Evaluating the evolutionary trees, constructed through every alignment on basis of all the deployed substitution models, the most-accurate tree is defined to be the one with the lowest log-likelihood score. The mutually synchronous behavior of the other two scores is subsequently evaluated for a considered set of alignment and substitution model, as an attempt to showcase the current pitfalls of these measures and to screen a more accurate assessment protocol for all the datasets. The evolutionary trees are lastly visualized through IToL (https://itol.embl.de/) for rationalizing the differences across various subfamilies (Sagulenko et al. 2018).

## Results and Discussion

### Building the sequence dataset

Despite purging the redundant hits, the datasets 1-3 orderly encompass 36, 109 and 712 sequences within the length range of 96-484, 84-4427 and 50-570 and encode a total of 6807, 150301 and 827623 residues respectively. It substantially increases the sequence space of these sets and only a few algorithms are found capable of building their alignments.

Unfortunately, no substitution matrix consistently yields the most accurate and biologically reasonable alignments for all sequence datasets (Kemena and Notredame 2009), and as an attempt to draw a more accurate evolutionary relationship, most of the phylogeny protocols use the bootstrap methodology (Huson and Bryant 2006). Although presently taken to be key score for finding the best evolutionary model for a sequence dataset, the log-likelihood score may not always select the most-accurate tree always and it should be strengthened with added measures to make the analysis more accurate. An evolutionarily closer sequence, often sharing a higher sequence identity, should have a smaller branch length, and this should strengthen the approximation of the log-likelihood metric (Parks and Goldman 2014). While the datasets 1, 2 and 3 show a sequence identity within the range of 0-99.52, 0-97.97 and 0-99.53 respectively, the average sequence identity is orderly found to be of 27.452±21.586, 16.742±7.815 and 29.137±12.759, and it suggests a higher evolutionary divergence among the set2 sequences.

### Evolutionary analysis for three datasets

The maximum-likelihood based phylogenetic tree is constructed through the ultrafast bootstrap methodology of IQTree to save computational resources and time (Nguyen et al. 2015). The number of free parameters (*branch + model parameters*) for the three sets in all the alignments is orderly found to be 88, 234, 1440. For the defined sequence sets and all of their constructed trees, the assessment is done through the log-likelihood score (Chatzou et al. 2016; Parks and Goldman 2014), total tree length (Nguyen et al. 2015), and the sum of the internal branch lengths (Chang et al. 2014).

Correlating the maximum, minimum, average and standard deviation scores for the 13 constructed Clustal-OMEGA, MUSCLE, KALIGN, MAFFT, TCOFFEE1, ESPRESSO TCOFFEE, MCOFFEE, PSI TCOFFEE, MUSCLE1, Clustal1, MAFFT1, KALIGN1 and PROMALS3D alignments against the 9 substitution models, viz. BLOSUM62, Dayhoff, JTT, JTT DCMut, LG, PMB, VT, WAG, DCMut for set1, it is observed that the total tree length and total internal branch length follow a similar scoring pattern, as expected (Figure 2). However, the scoring undulation for the log-likelihood score is found quite different, and it falls in line with the recent publication (Sievers and Higgins 2018). A similar trend is observed for the set2 and set3 datasets. The residue substitution scores are plotted for the constructed alignments (Figure 3), wherein the X and Y axis orderly refers to substitution matrices and alignment protocols. For sets1-3, the total internal branch length and total tree length are found lowest for BLOSUM62 for the Clustal alignment in contrast to the highest respective scores shown by DCMut matrix for the PROMALS3D alignment. It indicates that the structure-based sequence alignment should not be directly deployed for a study, unless explicitly evaluated for the selected dataset. Hence, to derive a more accurate evolutionary relationship, the biologically correct sequence alignment and residue substitution model should be used (Pearson 2013a).

**Figure 2:**
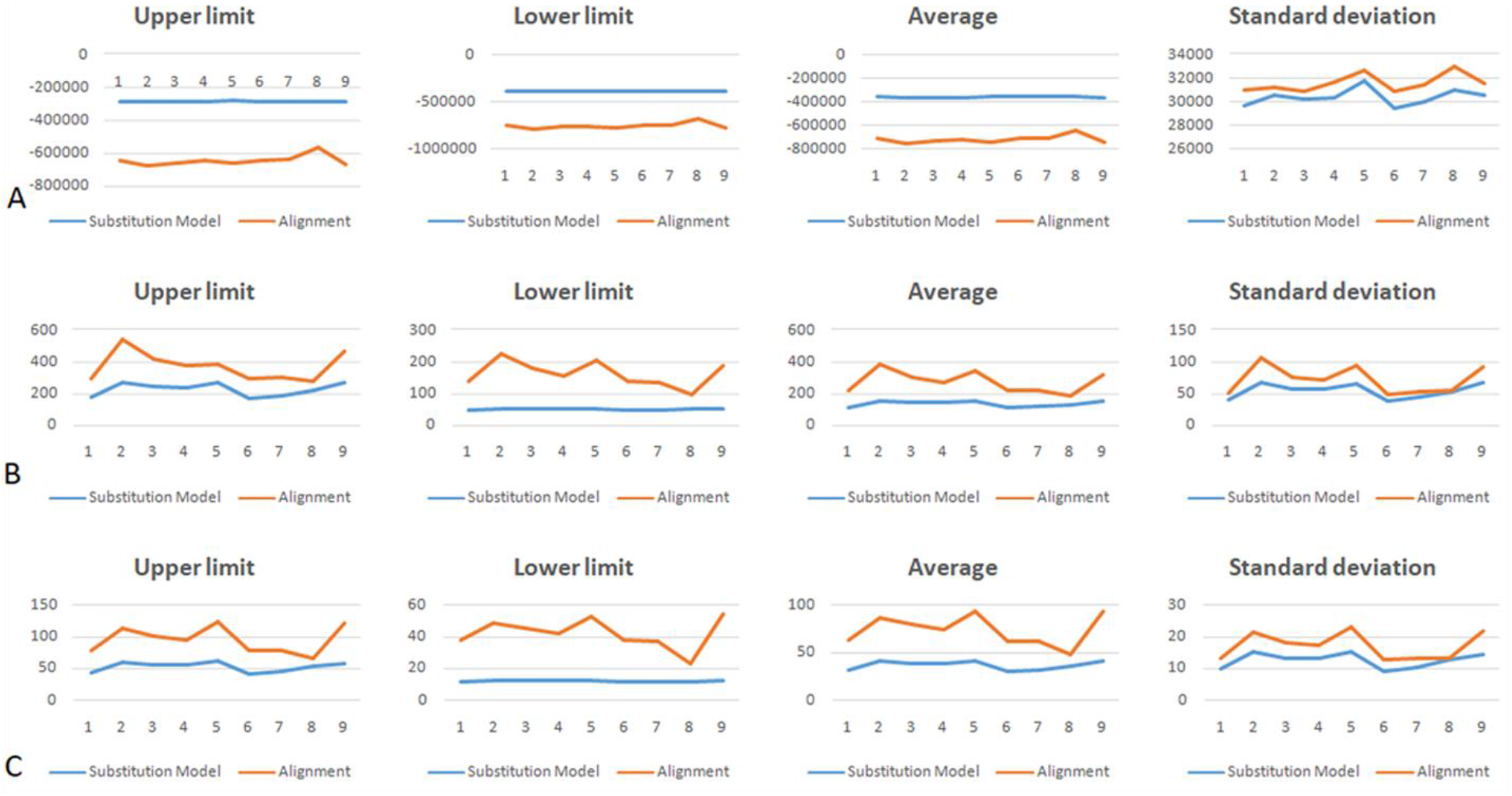
Maximum, minimum, average and standard deviation scoring undulations for (A) Log-likelihood score (B) total tree length (C) total internal branch length for the deployed substitution models (blue) and alignments (red) for the 36-sequence set1.

**Figure 3.**
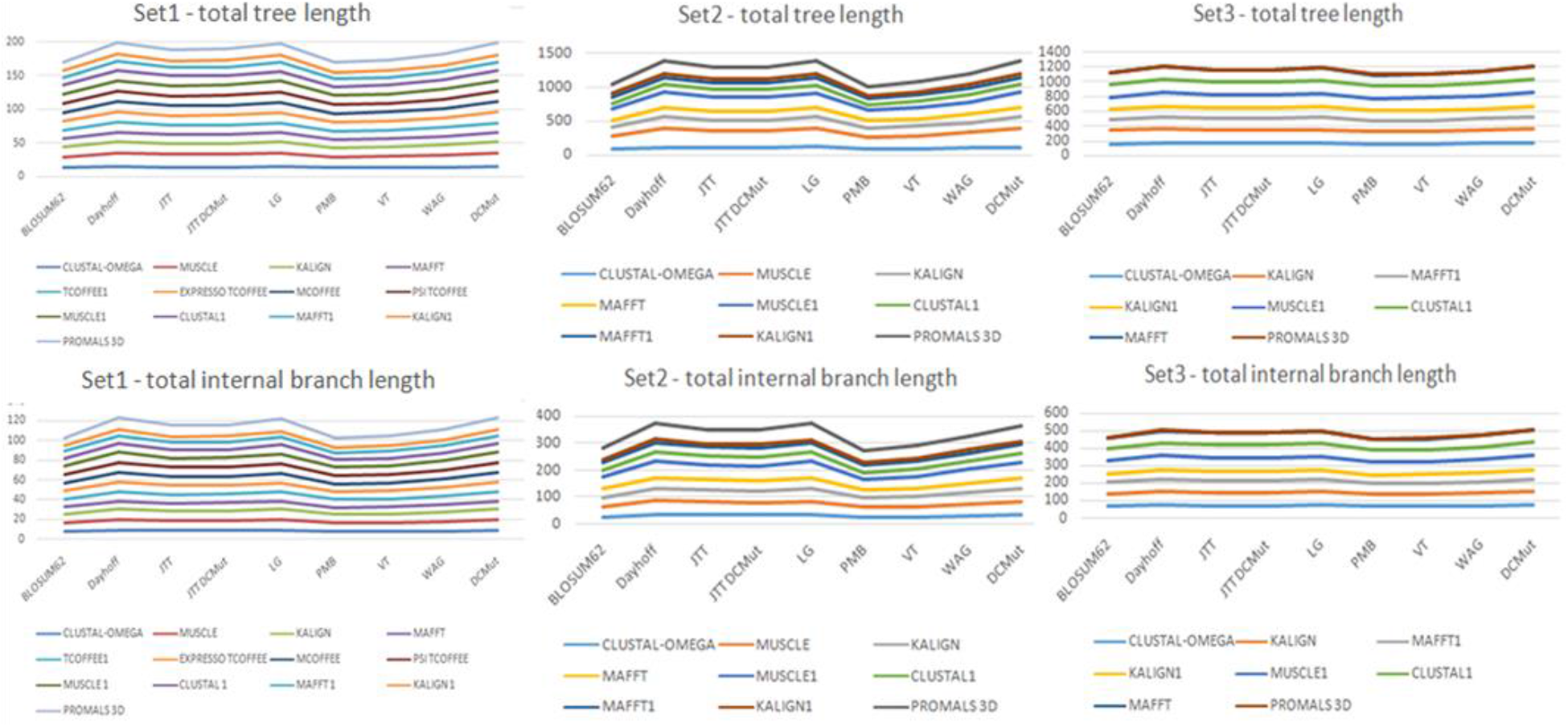
Scoring alterations of the total tree length and internal branch length of the three sequence datasets set1-set3 for the phylogenetic trees constructed through the alignments on the basis of the selected substitution models.

### Finding the top-ranked alignments and substitution models

For a protein sequence dataset, the evolutionary trees are drawn on the basis of 9 substitution matrices and 13 sequence alignments for sets 1-3 respectively, and assessed through the log-likelihood, total tree-length and total internal branch-length scores. The evolutionary likelihood of a phylogenetic tree is majorly assessed through three scoring measures viz. log-likelihood (Nikoh et al. 1994), total tree length and total internal tree length (Nikoh et al. 1994), and a solution with the lowest score is expected to be an optimal one. Among the 279 evolutionary trees, the top-ranked phylogenetic solutions, yielded by the best alignments (Table I) and the more accurate substitution models (Table II), are evaluated for the lower and upper limits of the three scores. The parameters are likewise evaluated for the overall average scores for each alignment protocol and substitution model, and the consistently correct alignment (Table III) and the substitution model (Table IV) are screened.

**Table 1:** Lower and upper limit of the log-likelihood, total tree length and total internal tree length scores for the evolutionary tree of the top-ranked alignments, screened from the pool built through various substitution models of the three datasets. A lower score indicates a better solution.

|  |  | **Log-likelihood**<br>**Alignment protocol**<br>**(Substitution model)** | **Total tree length**<br>**Alignment protocol**<br>**(Substitution model)** | **Total internal tree length**<br>**Alignment protocol**<br>**(Substitution model)** |
| --- | --- | --- | --- | --- |
| **Set1** | **Lowest** | -11898.259<br>MUSCLE(LG) | 9.453<br>Kalign1 (PMB) | 5.289<br>Kalign1 (PMB) |
|  | **Highest** | -9982.515<br>Kalign1 (WAG) | 20.227<br>MUSCLE (LG) | 12.423<br>PROMALS3D (Dayhoff) |
| **Set2** | **Lowest** | -391294.242<br>MUSCLE (Dayhoff) | 48.06<br>Kalign1 (PMB) | 11.301<br>Kalign1(PMB) |
|  | **Highest** | -281093.705<br>Kalign1 (LG) | 272.449<br>MUSCLE (DCMut) | 62.593<br>PROMALS3D (LG) |
| **Set3** | **Lowest** | -128789.188<br>PROMALS3D (LG) | 129.367<br>Kalign1 (VT) | 51.48<br>Kalign1 (PMB) |
|  | **Highest** | -120287.5159<br>MUSCLE1 (JTT) | 202.405<br>PROMALS3D (Dayhoff) | 84.51<br>PROMALS3D (DCMut) |

**Table 2:** Lower and upper limit of the log-likelihood, total tree length and total internal tree length scores for the evolutionary tree constructed through the top-ranked substitution models for the three datasets. A lower score indicates a better solution.

|  |  | **Log-likelihood**<br>**Best substitution model** | **Total tree length**<br>**Best substitution model** | **Total internal tree length**<br>**Best substitution model** |
| --- | --- | --- | --- | --- |
| **Set1** | **Lowest** | -11898.259<br>MUSCLE(LG) | 9.453<br>Kalign1 (PMB) | 5.289<br>Kalign1 (PMB) |
|  | **Highest** | -9982.515<br>Kalign1 (WAG) | 20.227<br>MUSCLE (LG) | 12.423<br>PROMALS3D (Dayhoff) |
| **Set2** | **Lowest** | -391294.242<br>MUSCLE (Dayhoff) | 48.06<br>Kalign1 (PMB) | 11.301<br>Kalign1(PMB) |
|  | **Highest** | -281093.705<br>Kalign1 (LG) | 272.449<br>MUSCLE (DCMut) | 62.593<br>PROMALS3D (LG) |
| **Set3** | **Lowest** | -128789.188<br>PROMALS3D (LG) | 129.367<br>Kalign1 (VT) | 51.48<br>Kalign1 (PMB) |
|  | **Highest** | -120287.5159<br>MUSCLE1 (JTT) | 202.405<br>PROMALS3D (Dayhoff) | 84.51<br>PROMALS3D (DCMut) |

**Table 3:**
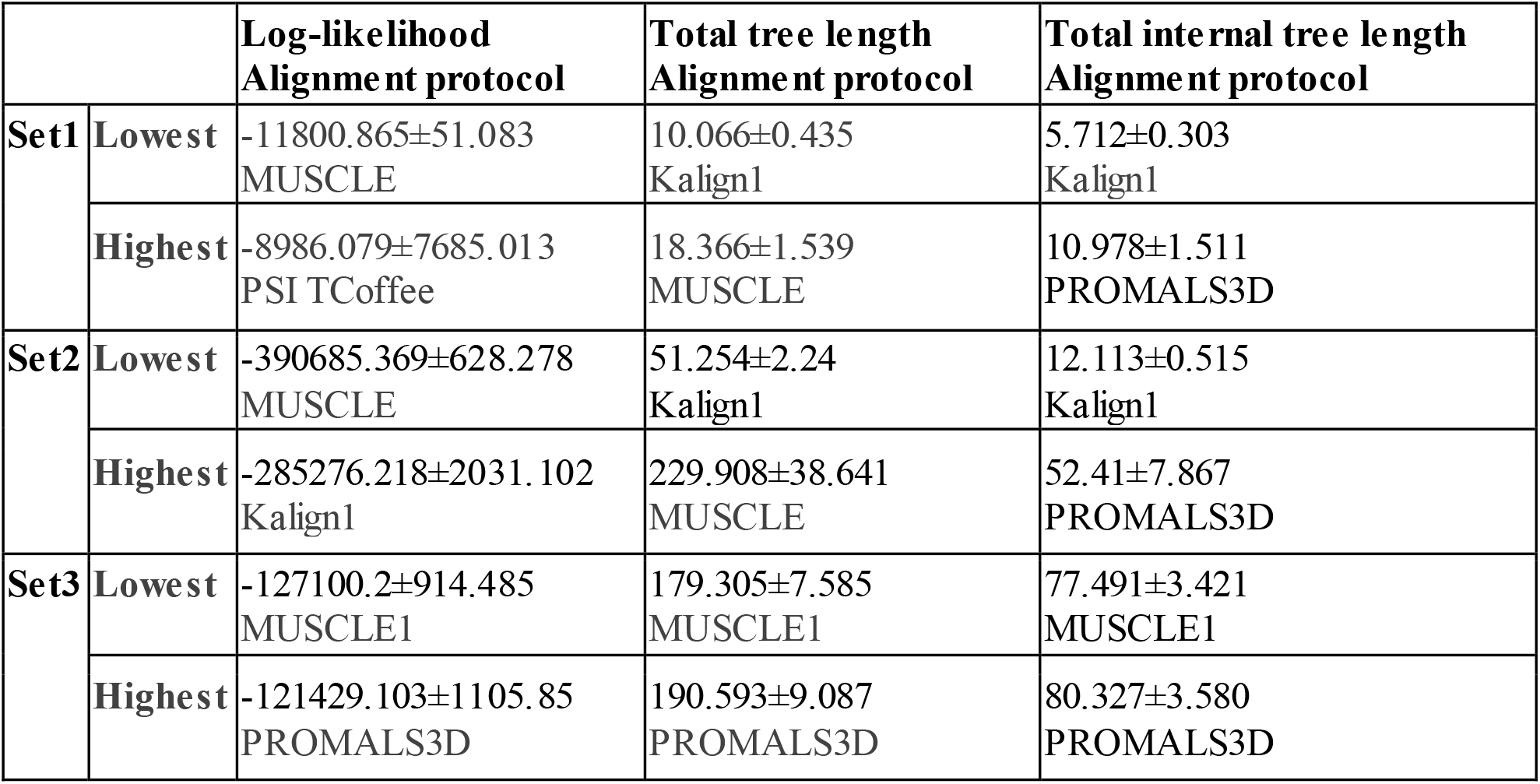
Lowest and highest average score of the log-likelihood, total tree length and total internal tree length scores for the evolutionary tree top-ranked alignments of the three datasets. A lower score indicates a better solution.

**Table 4:** Lowest and highest average score of the log-likelihood, total tree length and total internal tree length scores for the evolutionary tree constructed through the top-ranked substitution models for the three datasets. A lower score indicates a better solution.

|  |  | **Log-likelihood**<br>**Best substitution model** | **Total tree length**<br>**Best substitution**<br>**model** | **Total internal tree length**<br>**Best substitution model** |
| --- | --- | --- | --- | --- |
| **Set1** | **Lowest** | -11376.241±487.047<br>LG | 13.072±1.637<br>PMB | 7.861±0.973<br>BLOSUM62 |
|  | **Highest** | -9474.36±6320.371<br>WAG | 15.418±2.271<br>Dayhoff | 9.513±1.54<br>Dayhoff |
| **Set2** | **Lowest** | -358941.659±29511.738<br>Dayhoff | 112.637±39.248<br>PMB | 30.21±9.284<br>PMB |
|  | **Highest** | -357079.167±31845.036<br>WAG | 155.0811±67.738<br>LG | 41.579±15.52<br>Dayhoff |
| **Set3** | **Lowest** | -120369.906±4838.96<br>LG | 159.124±14.712<br>PMB | 65.974±7.549<br>PMB |
|  | **Highest** | -117866.669±5195.16<br>JTT | 175.996±18.798<br>Dayhoff | 70.992±8.940<br>JTTDCMut |

Ranking the evolutionary trees to find the top-ranked alignment protocols for the three sets, it is observed that the MUSCLE, MUSCLE and PROMALS3D show the lowest log-likelihood score of −11898.259, −391294.242 and −128789.18 against the respective scores of −9982.515, −281093.705 and −120287.51 for the Kalign1, Kalign1 and MUSCLE1 alignments. Assessing with the minimal tree length and internal tree length measure, Kalign1, Kalign1 and PROMALS3D and Kalign1, Kalign1 and MUSCLE1 alignments are orderly found to construct better trees for sets1-3, against the respective solutions of MUSCLE, MUSCLE and MUSCLE1, and PROMALS3D, PROMALS3D and PROMALS3D (Edgar 2004; Pei et al. 2008). For the 9 low-scoring evolutionary trees for sets 1-3 for all the scoring measures, KAlign1, MUSCLE, MUSCLE1 and PROMALS3D are orderly found to yield the best solutions for 4, 2, 2 and 1 cases, and PMB, LG, JTT, Dayhoff models are found to yield the best solution for 5,1, 1, 1 and 1 cases (Arenas 2015; Pei and Grishin 2014).

Evaluating the set1-set3 trees with log-likelihood and as per the substitution models, LG, Dayhoff and BLOSUM62 are found to yield more accurate solutions through MUSCLE, MUSCLE and PROMALS3D alignments, against the respective worst trees formed by VT, LG and JTT models for the Kalign1 alignments (Pei and Grishin 2014). For the lowest tree length and internal tree length scores for sets1-3, Kalign1 consistently yields the best trees through PMB, PMB and BLOSUM62 matrices, against the respective worst trees formed by LG, DCMut and Dayhoff, and Dayhoff, DCMut and Dayhoff substitution models through MUSCLE, MUSCLE and PROMALS3D, and PROMALS3D, MUSCLE and MUSCLE1 alignments. However, screening the top-ranked 9 trees on the basis of the substitution models, PMB, BLOSUM62, LG, Dayhoff, DCMut based solutions are found the best for 4, 2, 1 and 1 cases, although 7 KAlign1 and 2 MUSCLE alignments are only found to build these 9 trees. Amongst the 18 evolutionary trees constructed for various sequence alignments and substitution models of the three sets, MUSCLE, MUSCLE1, Kalign1 and PROMALS3D protocols are observed to consistently yield more accurate trees for 4, 1, 11 and 2 cases for various substitution models viz. LG, PMB, Dayhoff, JTT and BLOSUM62. Hence, the alignment and substitution matrix protocols are statistically evaluated to find the consistently correct evaluation measure (Pei and Grishin 2014). Screening the best alignment protocols on the basis of the average statistics of the three measures, Kalign1, MUSCLE1 and MUSCLE protocols are found to yield the best trees for the 4, 3 and 2 cases, and MUSCLE1 yields the consistently correct evolutionary relationships for set3 for all scoring parameters. However, for sets 1 and 2, MUSCLE is found to yield the best log-likelihood trees and Kalign1 is found to build the trees with the lowest total tree length and internal tree length scores. Further, the standard deviation score for these three protocols is respectively found within the percentage range of 4.251 - 5.304. Conversely, the Kalign1 alignment is found to show the worst tree with the highest log-likelihood score for VT, LG and BLOSUM62 matrices for sets 1-3, and likewise is the case for MUSCLE and PROMALS3D for the tree-length based scores.

A similar analysis of the substitution measures likewise unfolds another complicacy. It is observed that PMB, LG, Dayhoff and BLOSUM62 yield the best trees for 5, 2, 1 and 1 cases, with PMB model consistently being more accurate for the tree-length based measure. As per the total tree-length measure, the PMB matrix is found to be the best for all the three datasets (Philippe et al. 2011; Shalini Veerassamy 2003). For the internal tree length, BLOSUM62 and PMB matrix yield the best tree for set1 and sets 2 and 3, and for log-likelihood, LG and Dayhoff models aid the construction of sets1 and 3, and set2 respectively. Further, the standard deviation against mean is found within the percentage range of 4.02 - 34.84 for the LG and PMB matrices for the log-likelihood and total tree length score for sets 3 and 2 respectively.

The analysis proves that both alignment protocol and substitution matrix are equally responsible for the construction of a biologically meaningful evolutionary tree (Jordan and Goldman 2012), and it indicates that the accuracy of an evolutionary protocol is highly dependent on the correct combination of these two protocols (Chang et al. 2014). Hence, the huge set of 270 (set1-117, set2-81, and set3-72) phylogenetic trees are reanalyzed to decipher the top-ranked pairs of sequence alignment and the substitution models, and to find their consistently correct algorithmic set (Table V).

**Table 5:**
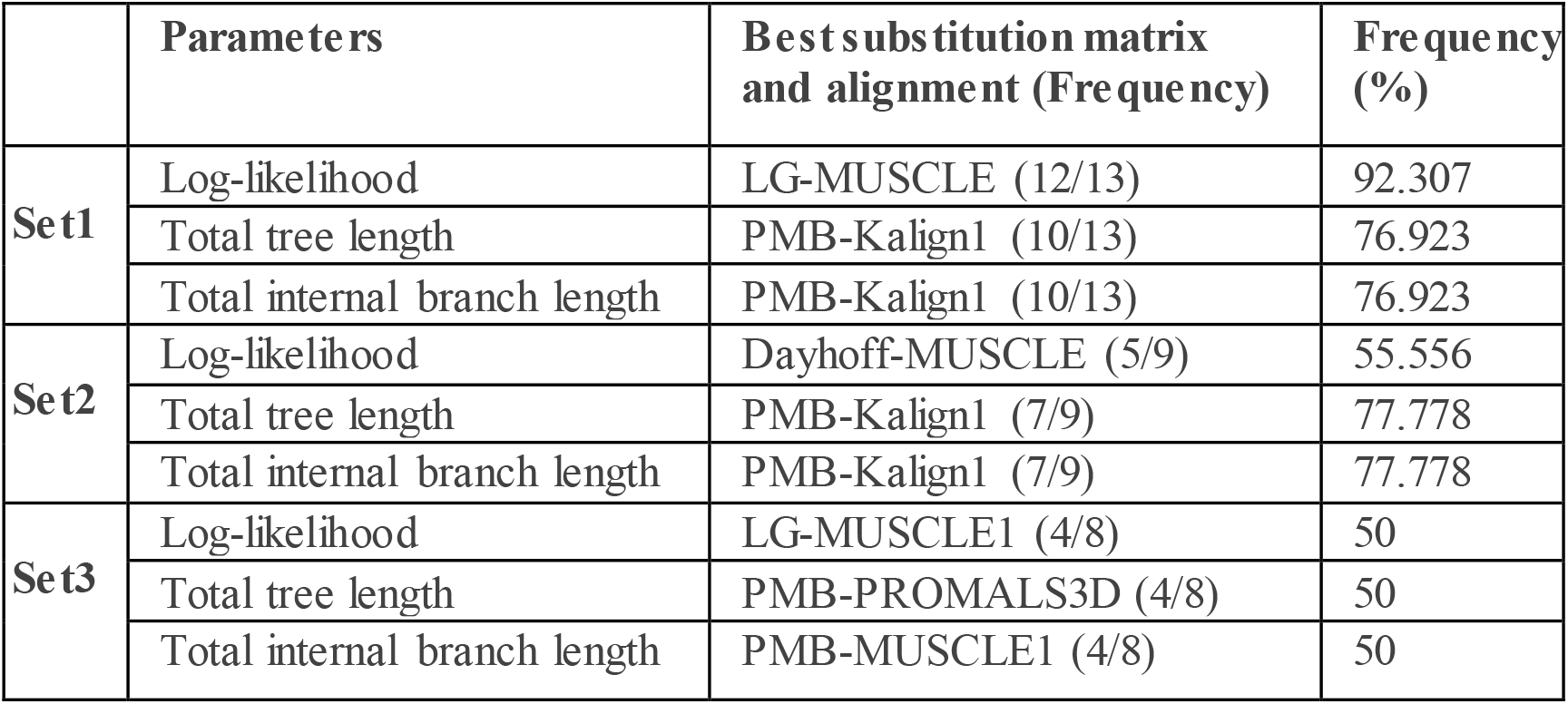
Top-ranked alignment-substitution model protocols for the three datasets for the three assessment measures: Log-likelihood, total tree length and total internal branch length. The failure to yield the consistently correct evolutionary model for any sequence dataset is indicated.

The analysis unleashes several intriguing features. MUSCLE alignment is found successful only for the smaller sequence datasets 1 and 2 (Deorowicz et al. 2016). The MUSCLE alignment, along with the LG and Dayhoff matrices, is found to yield the tree with the lowest log-likelihood score for 92.307% and 55.556% cases for the sets 1 and 2 respectively. For 270 constructed trees (92.307%), the MUSCLE alignment is found to work well with only the LG and Dayhoff substitution matrices for sequences shorter than 500 residues for the log-likelihood measure (Pearson 2013b), although for the distance based measure, the PMB matrix yields a more accurate tree through the -Kalign1 alignment (Nuin et al. 2006). However, the evolutionary trees constructed through it are maximally found better for the log-likelihood measure only, and its application is therefore not well-suited for drawing a robustly accurate phylogenetic relationship in view of the rapidly growing sequence data (Salichos et al. 2014). Further, PMB-Kalign1 protocol is found to yield the tree with the lowest tree length scores for 10 of the 13 set1 trees (76.923%) and 7 of the 9 set2 (77.778%) trees. However, for 712-sequence set3, the PMB matrix yields the tree with the lowest tree-length score for the PROMALS3D and MUSCLE1 alignments for set2 and set3 respectively. Although, PMB substitution matrix is found to consistently construct more accurate trees with the lowest total tree length score for all the sequence datasets, it is found incompatible to handle the diverse datasets with varying size and length (Stamatakis 2014). A substantial divergence is observed in the tree-topology of the top-ranked tree against the sub-optimal solutions (Shukla et al. 2019). It implies that there is a specific biological upper limit for each alignment protocol to derive the meaningful results of the overlapping residues, and is in line with the earlier study (C S Troy 2001). Evaluating the trees for the location of various branches, it could be concluded that an accurate alignment is mandatory to draw the best-possible evolutionary relationship, as has been recently shown (Georgios A Pavlopoulos 2010). A significant disparity is found for all the trees for all the scoring parameters (Hall 2013), and selection of the best tree is further hereby shown to be difficult, especially for bigger dataset (Nguyen et al. 2015).

As an attempt to build more accurate evolutionary trees, the protocols including RaxML iteratively construct the alignments with a guide tree (Stamatakis 2014). Further, the “phylogeny-aware” tools like PRANK (Ari Löytynoja 2005) and PAGAN (Loytynoja et al. 2012) presume such guide-trees as the true trees to ensure that the constructed alignments are evolutionarily correct, and are thus termed as the post-tree analysis methods. Although the interdependence of MSA and inference of a phylogeny tree has been described in these methods, the best combination of alignment method and substitution matrix has not been greatly excavated for assessments.

### Required strategic improvement over the existing protocols

The sequence alignments are usually constructed through several methodologies, and are substantially different for the number and placement of gaps, and as incorrect gaps lead to an inaccurate evolutionary tree, the alignments have always been iteratively constructed to find the best possible alignment and to derive the optimal phylogenetic solution (Blackburne and Whelan 2013; Fernández-Baca et al. 2004; Landan and Graur 2007; Sievers et al. 2013; T Heath Ogden 2006). Likewise, the substation models are usually deployed to derive the evolutionary relationship among proteins, and different models have been tried for constructing a more accurate tree (Duchene et al. 2016; Hoff et al. 2016; Shapiro et al. 2006). However, for a sequence dataset, the phylogeny servers never iteratively optimize the evolutionary tree to save their processing time and computational resource. As demonstrated by our study, the biological significance of alignments/substitution models should be simultaneously tested through iterative assessments to construct the best evolutionary model for a sequence dataset. The study showcases a strategy to circumvent the potential pitfalls of the evolutionary algorithm. It will allow us consistently construct an evolutionary correct model for a sequence dataset and will subsequently improve the prediction accuracy of several different types of biological studies, be it a sequence characterization (Guder and Krishna 2019; Kalyani et al. 2019; Mallu et al. 2019; Satyanarayana et al. 2017; Somavarapu et al. 2017; Vemuri et al. 2019) or proteomic study (Jaswanthi et al. 2019; Rao et al. 2017; Samara Shekar Reddy et al. 2019; Srideepthi et al. 2017).

### Strategic steps to assess and achieve biological accuracy of alignment and phylogeny

As recently shown by an evolutionary analysis of the foldIV transaminases with experimentally solved structures (Pavkov-Keller et al. 2016), all the four subfamilies, viz. l-branched chain aminotransferases (BCATs), d-amino acid aminotransferases (D-ATAs), (R)-selective transaminases (RATA) and 4-amino-4-deoxychorismate lyases (ADCL), are found to be clustered into different clades (Francesco G. Mutti 2011; Percudani and Peracchi 2009). However, the phylogenetic protocol failed to accurately classify two functionally characterized RATA sequences CpuTA1 and MgiTA1, sharing a 49% sequence identity and containing conserved amino acids for both RATA and ADCL (Pavkov-Keller et al. 2016). It thus shows the need of functional investigation of the input sequences before constructing their biologically meaningful tree. Accurate establishment and quantification of homology through various substitution model/alignment algorithms is proven to be the key feature to accurately align the sequences and generate the meaningful phylogenetic trees. Moreover, evolutionary biologists should thus construct the consensus tree, as implemented by IQ-Tree server (Nguyen et al. 2015), and investigate the chance of shrouded negative factors through iterative assessments.

## Conclusion

Evolutionary trees have long been deployed for several research methodologies. Here, we have focused on constructing an improved phylogenetic tree for a sequence dataset. The constructed set of 270 (set1-117, set2-81, set3-72) phylogenetic trees for the 36-sequence, 109-sequence and 712-sequence datasets illustrates that the accuracy of an evolutionary study is significantly determined by the compatibility of the sequence alignment and the residue substitution model, and is not simply governed by the usually parameterized bootstrap and minimum correlation coefficient features. The best complementary set of biologically closest alignment and the substitution model is observed to yield the most-accurate evolutionary tree. Assessing the resultant phylogeny solutions on the basis of log-likelihood and minimal internal branch length, the study streamlines the methodology and opens avenues to design the robustly accurate phylogeny protocol. The study also lays a benchmark for the development of a protocol to synergistically deploy the assessment measures and consistently derive a more accurate evolutionary solution for all datasets.

